# Early flowering in oilseed-type *Brassica rapa* plants results from nonsense-mediated mRNA decay (NMD) of *BrFLC2*

**DOI:** 10.1101/2021.05.05.442798

**Authors:** Sujeong Kim, Jin A Kim, Hajeong Kang, Dong-Hwan Kim

## Abstract

Many *Brassica* species require vernalization (long-term winter-like cooling) for transition to the reproductive stage. In the past several decades, scientific efforts have been made to discern the molecular mechanisms underlying vernalization in many species. Thus, to identify the key regulators required for vernalization in *Brassica rapa* L., we constructed a linkage map composed of 7,833 single nucleotide polymorphism (SNP) markers using the late-flowering Chinese cabbage (*B. rapa* L. ssp. *pekinensis*) inbred line ‘Chiifu’ and the early-flowering yellow sarson (*B. rapa* L. ssp. *trilocularis* (Roxb.)) line ‘LP08’ and identified a single major QTL on the upper-arm of the chromosome A02. In addition, we compared the transcriptomes of the lines ‘Chiifu’ and ‘LP08’ at five vernalization time points, including both non-vernalized and post-vernalization conditions. We observed that *BrFLC2* was significantly downregulated in the early flowering ‘LP08’ and had two deletion sites around the *BrFLC2* genomic region compared with the *BrFLC2* genomic region in ‘Chiifu.’ In the present study, we also demonstrate that early flowering in ‘LP08’ line is attributed to the low expression of *BrFLC2*, which is caused by nonsense-mediated mRNA decay (NMD). Therefore, this study provides a better understanding of the molecular mechanisms underlying floral transition in *B. rapa*.

**One sentence summary:** NMD-mediated degradation of *BrFLC2* mRNA transcripts is the main cause of rapid flowering of oilseed-type *B. rapa* ‘LP08’ plants.

## INTRODUCTION

Plants continually monitor environmental cues and optimize their growth and development. Vernalization, the long-term winter-like cold, plays an important role in floral transition in many biennial and perennial plants, including *Brassica rapa* L. (Amasino 2010). For decades, several efforts have been made to understand the molecular mechanisms underlying vernalization, particularly using the winter-annual *Arabidopsis* model plants. *A. thaliana* models can be divided into the following two groups depending on the flowering behavior: early flowering accessions referred to as summer annuals (for example, *Columbia-0*), which complete their life cycle in one growing season regardless of vernalization, and late flowering accessions called winter annuals (for example, *FRI_col*), which complete their life cycle in the next growing season after an intervening winter season (Kim et al. 2009).

Vernalization in the winter-annual Arabidopsis accessions requires a single locus named *FRIGIDA* (*FRI*) (Koornneef et al. 1994; Lee et al. 1994). *FRI* encodes a plant-specific nuclear protein, which substantially increases the expression of a potent floral repressor *FLOWERING LOCUS C* (*FLC*) (Geraldo et al. 2009). *FLC* codes for a MADS-box-containing DNA-binding protein, which suppresses a group of genes named as floral integrators, including *FLOWERING LOCUS T* (*FT*) and *SUPPRESSOR OF OVEREXPRESSION OF CONSTANS 1* (*SOC1*). High expression of *FLC* significantly impedes floral transition and delays flowering in winter-annual Arabidopsis (Dennis et al. 2006; Searle et al. 2006). Vernalization inhibits *FLC* transcription to release floral integrators *FT* and *SOC1* from the suppression of *FLC* (Sung and Amasino 2005). Thus, floral transition is accelerated by an increased expression of floral integrators following vernalization.

Chinese cabbage (*B. rapa* L. ssp. *pekinensis*), belonging to the family *Brassicaceae*, includes 37 species such as pak choi, cabbage, broccoli, radish, and turnip. These plant species are cultivated globally and are popular in Asian countries (Leijten et al. 2018). *B. rapa* includes both spring-type plants, which do not require vernalization and exhibit rapid flowering, and winter-type plants, which need vernalization for floral transition. Recently, several quantitative trait loci (QTL), affecting vernalization-mediated floral transition in *B. rapa*, were identified using independent QTL analyses. An RFLP-based genetic linkage map generated from the F_2_ population of a cross between an annual and a biennial oilseed cultivar identified two QTL peaks on chromosomes LG2 and LG8 (Teutonico and Osborn 1995; Osborn et al. 1997).

The reference genome of *B. rapa* inbred line ‘Chiifu-401-42’ (hereafter ‘Chiifu’) contains four *FLC* homologs, namely *BrFLC1–BrFLC3* and *BrFLC5. BrFLC1* and *BrFLC2* have been reported to be linked to the major QTLs controlling flowering time in Chinese cabbage (Li et al. 2009; Yuan et al. 2009). Additionally, a QTL analysis using early-flowering and late-flowering cultivars of *B. rapa* reported a major QTL linked to the *BrFLC2* region (Zhao et al. 2010; Wu et al. 2012; Xiao et al. 2013). In a separate study using the F_2_ population of a cross between the lines ‘Early’ × ‘Tsukena No. 2,’ QTL peaks were identified near *BrFLC2* and *BrFLC3* loci, wherein large insertions in the first intron of *BrFLC2* were found to hinder the vernalization-mediated stable repression of *BrFLC2*, delaying flowering in ‘Tsukena No. 2’ (Kitamoto et al. 2014).

The expression of all four *BrFLC* homologs decreases upon vernalization and stabilize at low levels even after exposure to warm temperatures (Kawanabe et al. 2016; Kim 2020). However, the molecular mechanisms underlying vernalization-mediated floral transition in Chinese cabbage is still not completely understood. Therefore, the aim of the present study was to identify the key regulators responsible for the flowering behavior of late-flowering Chinese cabbage ‘Chiifu’ and early-flowering yellow sarson ‘LP08’ lines. For this purpose, we performed SNP-based QTL mapping and comparative transcriptome analysis. For transcriptome analysis of vernalization response, RNA-seq data were generated and analyzed at five different time points, including non-vernalized (NV), 1-day cold (1V), 20-day cold (20V), 40-day cold (V40), and an after-vernalization (AV, 40-day cold followed by 10 days under warm conditions) treatments, for both the lines. We also investigated the role of nonsense-mediated mRNA decay (NMD) in vernalization in the two lines for better understanding of vernalization-mediated floral transition.

## RESULTS

### Flowering behavior in vegetable-type ‘Chiifu’ and oilseed-type ‘LP08’ *B. rapa*

Vegetable-type ‘Chiifu’ exhibited late flowering and required vernalization for bolting, while oilseed-type ‘LP08’ displayed early flowering without vernalization (Supplementary Fig. S1). To assess flowering behavior in ‘Chiifu’ and ‘LP08,’ the time required for bolting in the two lines were determined with (45-day vernalization treatment under long day (LD) conditions) and without vernalization. Under LD conditions without vernalization, the late-flowering ‘Chiifu’ did not flower even after approximately 120 d (counted as not-determined (ND)), while early-flowering ‘LP08’ took an average of 37 d to flower (DTF) (Fig. 1a). After the 45-d vernalization treatment, flowering in ‘Chiifu’ was accelerated, with an average of 23 DTF, indicating that this line requires vernalization for floral transition as previously described. However, the floral transition rate of ‘LP08’ after vernalization was relatively moderate, with an average of 17 DTF, suggesting that ‘LP08’ also responds to vernalization.

**Figure 1.**
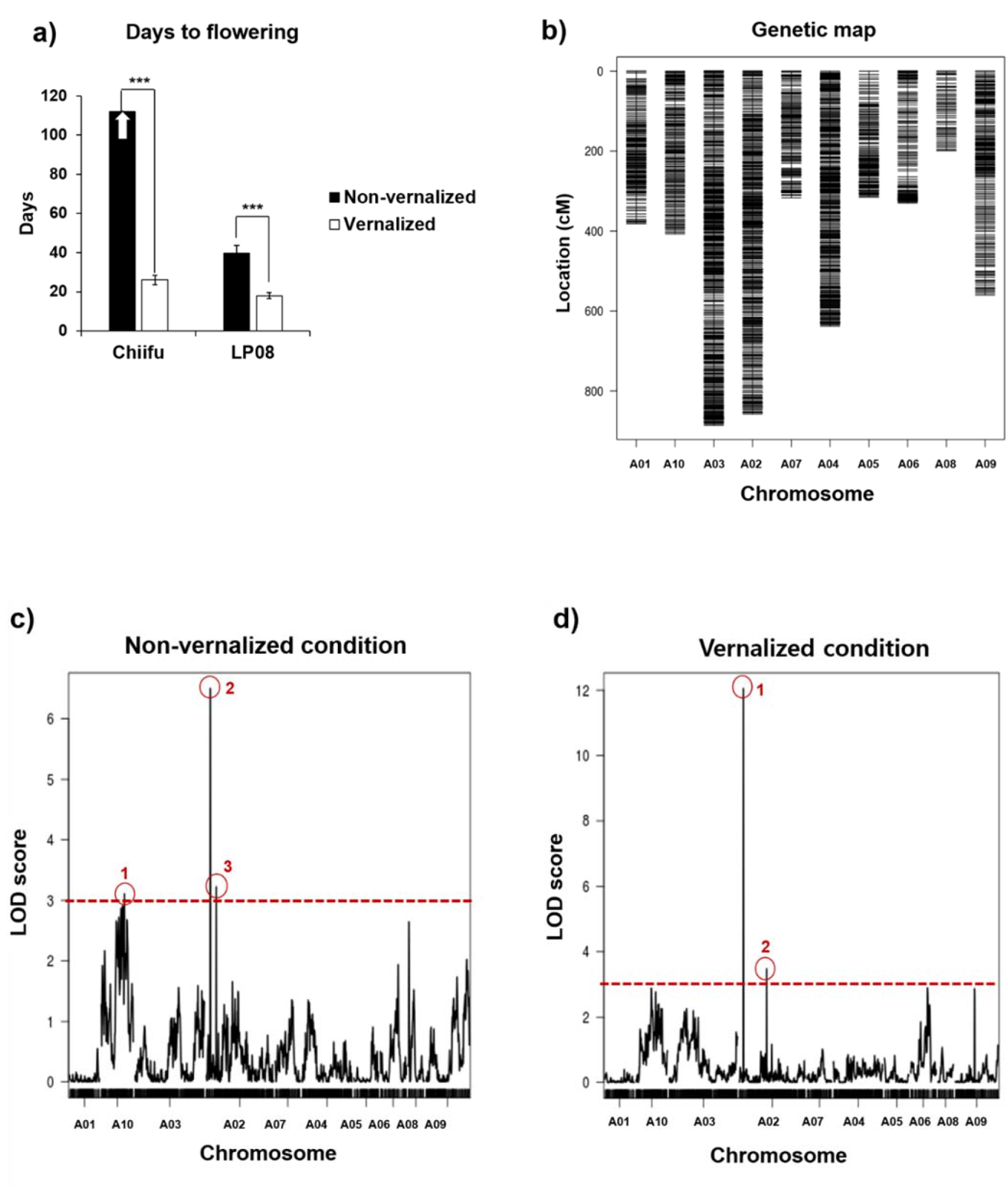
Construction of SNP-based linkage map and QTL mapping of flowering time. **(a)** Flowering time of late-flowering Chinese cabbage ‘Chiifu’ and rapid-flowering yellow sarson ‘LP08’ under both non-vernalized and vernalized conditions. **(b)** Linkage map generated using 7,833 single nucleotide polymorphisms (SNPs) identified between the two lines. **(c)** Results of QTL mapping of 151 F_5_ RILs to estimate the flowering time under non-vernalized condition. Significant peaks exceeding LOD score of 3 are indicated with red circle and numbers. **(d)** Results of QTL mapping of 151 F_5_ RIL lines to estimate the flowering time under vernalized (45-day cold) condition. Significant peaks exceeding LOD score of 3 are indicated with red circle and numbers.

### QTL mapping of flowering time

‘Chiifu’ and ‘LP08’ F_2_ lines were successively self-fertilized for five generations to produce a total of 151 recombinant inbred lines (RILs). Thereafter, a total of 96,506 SNPs were identified using the *B. rapa* reference genome and R/qtl package, and 7,833 SNPs were filtered using variant filtering. Only uniquely aligned reads were used to identify the genotypes of the RILs. The R package ASMap was used to determine the order of markers to construct linkage groups. The genetic linkage maps, which displayed 10 linkage groups corresponding to the 10 chromosomes of *B. rapa* (Fig. 1b), were used for QTL mapping of flowering time loci. After QTL mapping, peaks with LOD scores >3 were arbitrarily set as significant regions. Under non-vernalized (NV) condition, three QTL, with LOD scores >2, were detected on two chromosomes, A10 and A02 (Fig. 1c). Among the three QTL, the highest peak (>6 LOD score) was located on the upper arm of chromosome A02 (Fig. 1c). However, after vernalization, only two QTL, with LOD scores >3, were detected on chromosome A02. Similar to the NV treatment, the highest peak (>12 LOD score) during vernalization was also detected in the upper arm of chromosome A02 (Fig. 1d). Detection of the highest QTL peak in the same region of chromosome A02 (A02:1950553–A02:7744515) in both NV and vernalized conditions suggests that this region contains key regulators of flowering behavior in ‘Chiifu’ and ‘LP08’ lines.

### Comparative transcriptomic analysis of ‘Chiifu’ and ‘LP08’ lines after vernalization

To identify the candidate genes responsible for early flowering in ‘LP08’, we performed high-throughput RNA-seq analysis of ‘Chiifu’ and ‘LP08’ plants at five different vernalization time points, namely NV, 1-day cold treatment (V1), 20-day cold treatment (V20), 40-day cold treatment (V40), and 40-day cold treatment followed by 10 days under warm conditions (V40T10). A total of 30 RNA-seq libraries (including biological replicates) were constructed, sequenced, and analyzed, and correlation heatmap analyses revealed distinct clustered groups in the samples from each time point (Supplementary Fig. S1).

### Identification of candidate flowering genes in ‘Chiifu’ and ‘LP08’

QTL mapping revealed that a narrow region on the upper arm of chromosome A02 (approximately 5.8 kb, spanning A02:1950553–A02:7744515) contained major candidate genes responsible for rapid flowering in the ‘LP08’ line. We scanned the region and identified 228 annotated genes (Bra02000395–Bra02001489) (Supplementary Table S2), including five flowering-related genes (*BrFLC2, BrHY5a, BrEMF1a, BrVIN3a*, and *BrLD*).

Furthermore, to narrow down the list of causative genes for early flowering in ‘LP08,’ we compared the transcript levels of these flowering genes between ‘Chiifu’ and ‘LP08’ at all the vernalization time points. As shown in Fig. 2, *BrFLC2* was downregulated in the ‘LP08’ line at each time point (NV–V40T10), while it was highly upregulated in ‘Chiifu’ before vernalization (NV) and significantly downregulated after long-term cold treatments (V20 and V40). Moreover, the reduced level of *BrFLC2* in ‘Chiifu’ was stably maintained even after the plants were returned to a warmer temperature (V40T10), resembling the expression pattern of *Arabidopsis FLC* upon vernalization. Therefore, we hypothesized that the reduced expression of *BrFLC2* in ‘LP08’ markedly contributes to the rapid-flowering behavior in ‘LP08,’ because *BrFLC2* functions as a floral repressor.

**Figure 2.**
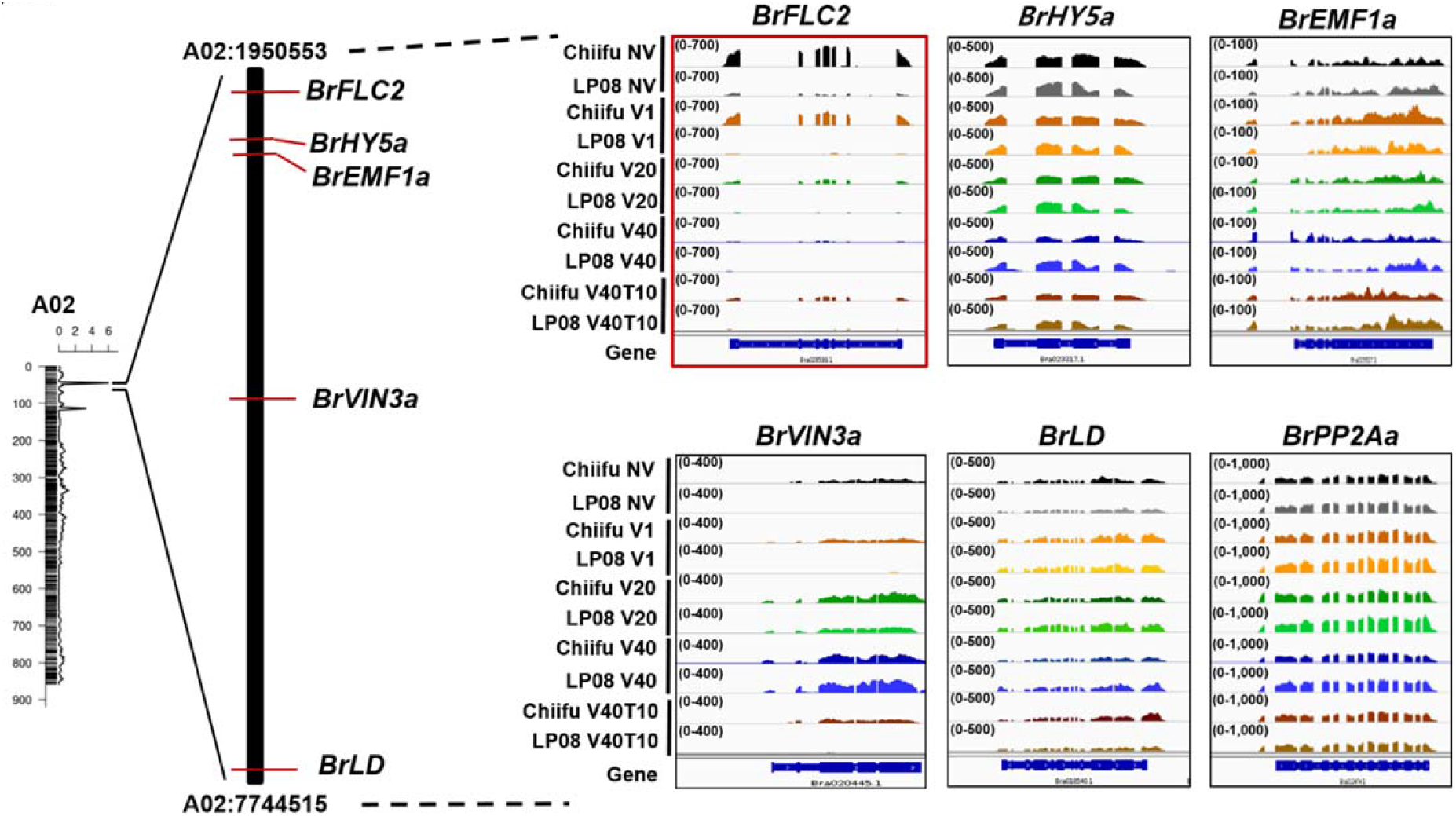
Transcript levels of candidate floral transition genes located on the upper arm of Chromosome A02. Integrative genome browser (IGV) illustration of the five flowering genes located on the upper arm of Chromosome A02 (A02:1950553–A02:7744515) showing a major QTL peak. *BrPP2Aa* was used as the reference gene to confirm the normalization of RNA-seq reads between ‘Chiifu’ and ‘LP08’ throughout the vernalization treatment (NV, V1, V20, V40, and V40T10). Different colors show transcript reads mapped to individual genes at different time points. Read coverage normalized by total number of mapped reads are indicated at the top left corner of each track in parentheses. NV: non-vernalized; V1: 1-day cold; V20: 20-day cold; V40: 40-day cold; V40T10: 40-day cold followed by 10 days under warm temperature.

We also compared the transcript levels of *BrHY5a, BrEMF1a, BrVIN3a*, and *BrLD* between the two lines. However, the transcript levels of these genes were similar between ‘Chiifu’ and ‘LP08’ at each vernalization time point, indicating that these flowering genes are not responsible for the early flowering behavior of ‘LP08’ (Fig. 2).

We also analyzed RNA-seq data for the four *FLC* homologs identified in ‘Chiifu’ and ‘LP08’ using *BrPP2Aa* (Bra012474) for normalization. Similar to the results shown in Figure 2, the transcript levels of *BrFLC2* were reduced in ‘LP08’ (Supplementary Fig. S2). In addition, we verified the transcript levels of the four *BrFLC* genes using qRT-PCR (Fig. 3a–3d) and again observed a similar *BrFLC2* expression pattern in the ‘LP08’ line compared with that of the ‘Chiifu’ line (Fig. 3b). Among other FLC homologs, *BrFLC1* and *BrFLC5* exhibited reduced expression in ‘LP08’ at NV and V1 time points (Fig. 3a and 3d), implying that low expression levels of *BrFLC1* and *BrFLC5* might also contribute to early flowering under NV conditions. However, ‘LP08’ had higher expression of *BrFLC1* and *BrFLC5* under long-term cold conditions (V20, V40, and V40T10) compared to their transcript levels in ‘Chiifu,’ indicating that they were not associated with vernalization-mediated acceleration of flowering time in ‘LP08.’ Moreover, the transcript levels of *BrFLC3* did not show a significant difference between ‘Chiifu’ and ‘LP08’ throughout vernalization treatment (Fig. 3c.

**Figure 3.**
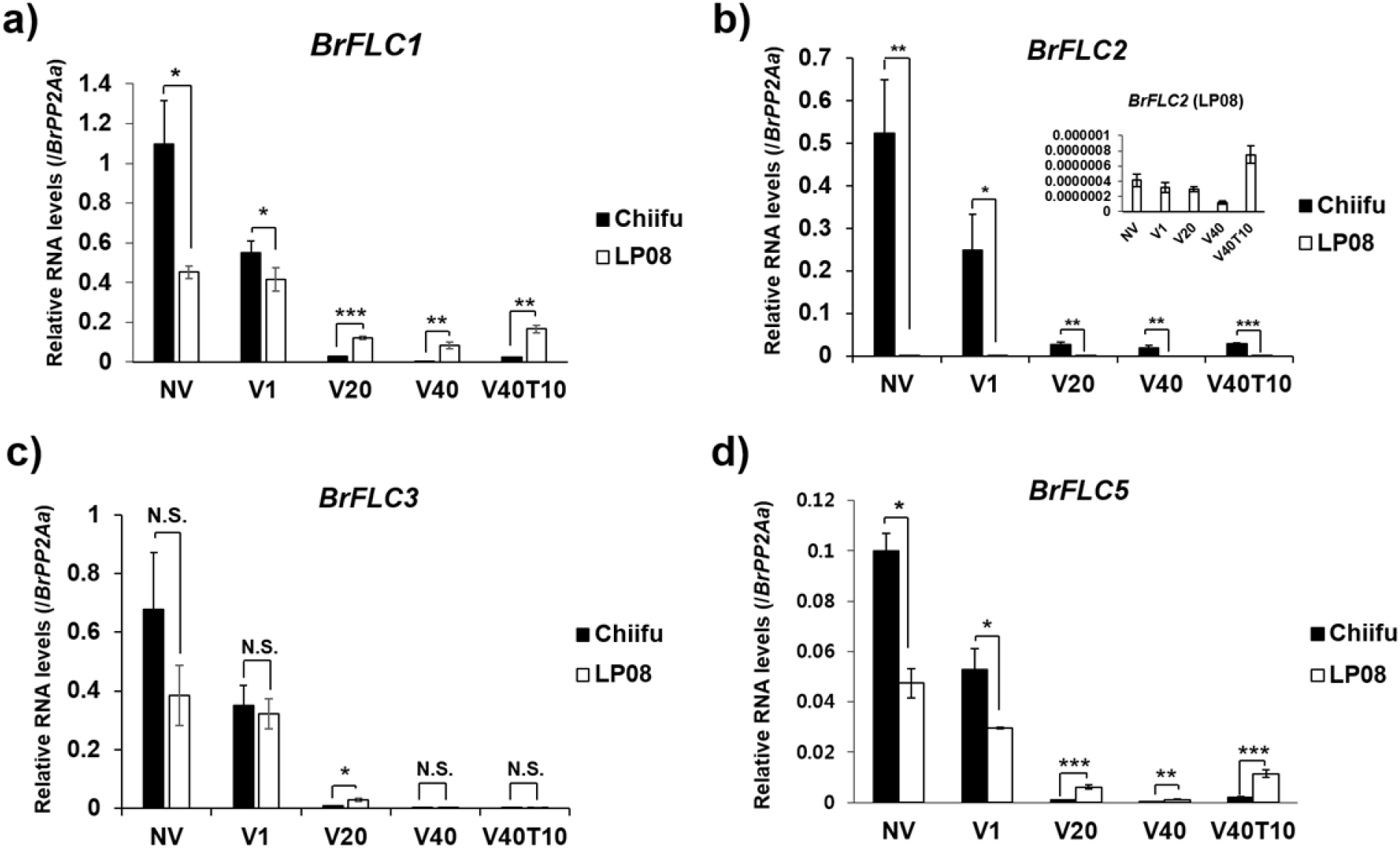
Transcript levels of the four *BrFLC* homologs in ‘Chiifu’ and ‘LP08’ along the vernalization time points. Results of qRT-PCR of the four *BrFLC* homologs, (**a**) *BrFLC1*, (**b**) *BrFLC2*, (**c**) *BrFLC3*, and (**d**) *BrFLC5*, throughout the vernalization treatment. In B, an inset indicates a magnified view of the levels of *BrFLC2* mRNA in ‘LP08’ along the vernalization time course. *BrPP2Aa* was used as the reference gene. Average and standard errors were calculated using the Ct values of three biological replicates. Significance was statistically determined using t-test (\**p* > 0.05; \*\**p* < 0.01; \*\*\**p* < 0.001).

### Transcription of *VIN3* homologs in ‘Chiifu’ and ‘LP08’

*AtVIN3* (*VERNALIZATION INSENSITIVE 3*), which encodes a PHD-finger protein, has been reported to play an important role in vernalization-mediated floral transition in *Arabidopsis*, along with *AtFLC* (Sung and Amasino 2004; Bastow et al. 2004). AtVIN3 directly binds to *FLC* and epigenetically suppresses the transcription of *AtFLC* during vernalization. Furthermore, *AtVIN3* is not expressed before vernalization but is upregulated during vernalization, with its expression decreasing rapidly after the plants are returned to warm temperatures. Therefore, we investigated the expression pattern of *VIN3* homologs in the two *B. rapa* lines.

Two homologs, *BrVIN3a* (Bra020445) and *BrVIN3b* (Bra006824), were identified in both ‘Chiifu’ and ‘LP08.’ The transcriptional pattern of *BrVIN3b* resembled that of *AtVIN3*, with reduced expression before vernalization (NV), high expression during vernalization (V1–V40), and severely reduced expression after vernalization (V40T10) (Fig. 4b). However, *BrVIN3a* exhibited a different transcription pattern in the ‘Chiifu’ line; the expression pattern of *BrVIN3a* in ‘LP08’ was similar to that of *AtVIN3* (Fig. 4a), while it was moderately expressed before vernalization (NV), had increased expression during vernalization (V20 and V40), and was moderately decreased after vernalization (V40T10) in ‘Chiifu’ (Fig. 5a). The fact that *BrVIN3a* is moderately expressed before and after vernalization in ‘Chiifu’ suggests that *BrVIN3a* plays additional roles during development other than those responsible for the vernalization response. However, *BrVIN3b* RNA-seq reads did not exactly match the predicted gene structure, with the first exon mapped to the promoter region of the gene (Supplementary Fig. S3), indicating that the identified version of *BrVIN3b* gene needed correction.

**Figure 4.**
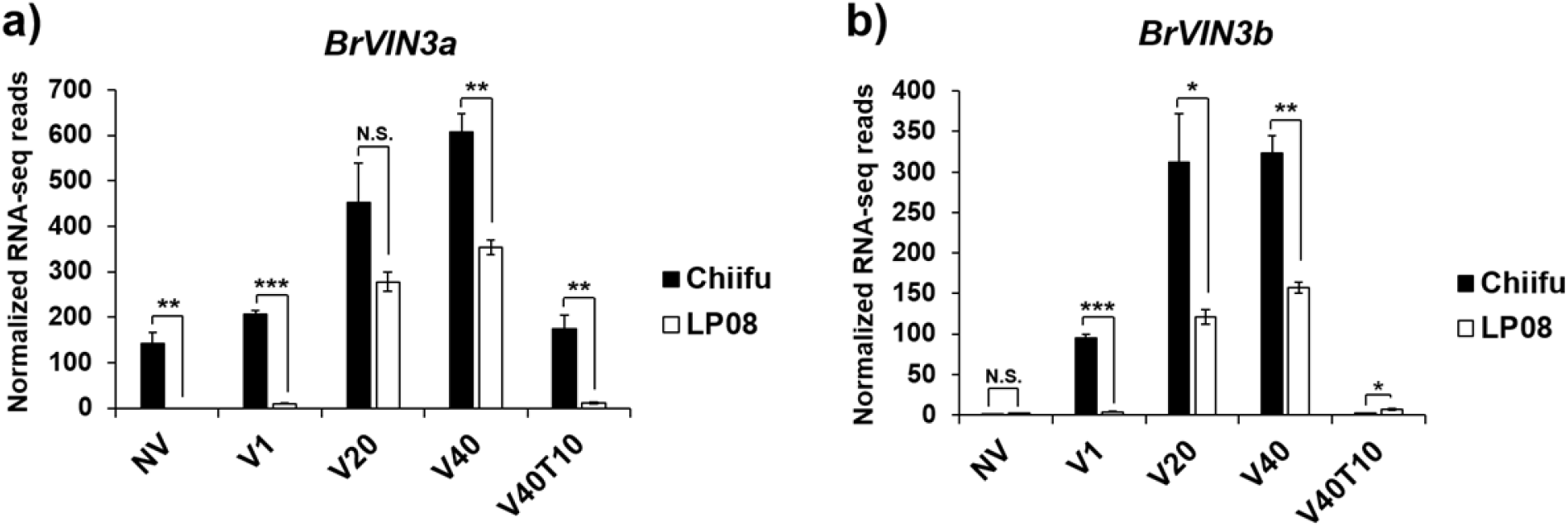
Transcript levels of the two *BrVIN3* homologs in ‘Chiifu’ and ‘LP08’ along the vernalization time points. Results of qRT-PCR of the two *BrVIN3* homologs, (**a**) *BrVIN3a* and (**b**) *BrVIN3b*, in ‘Chiifu’ and ‘LP08’ along the vernalization time points. *BrPP2Aa* was used as the reference gene. Average and standard errors were calculated using the Ct values of three biological replicates. Significance was statistically determined using t-test (\**p* > 0.05; \*\**p* < 0.01; \*\*\**p* < 0.001).

**Figure 5.**
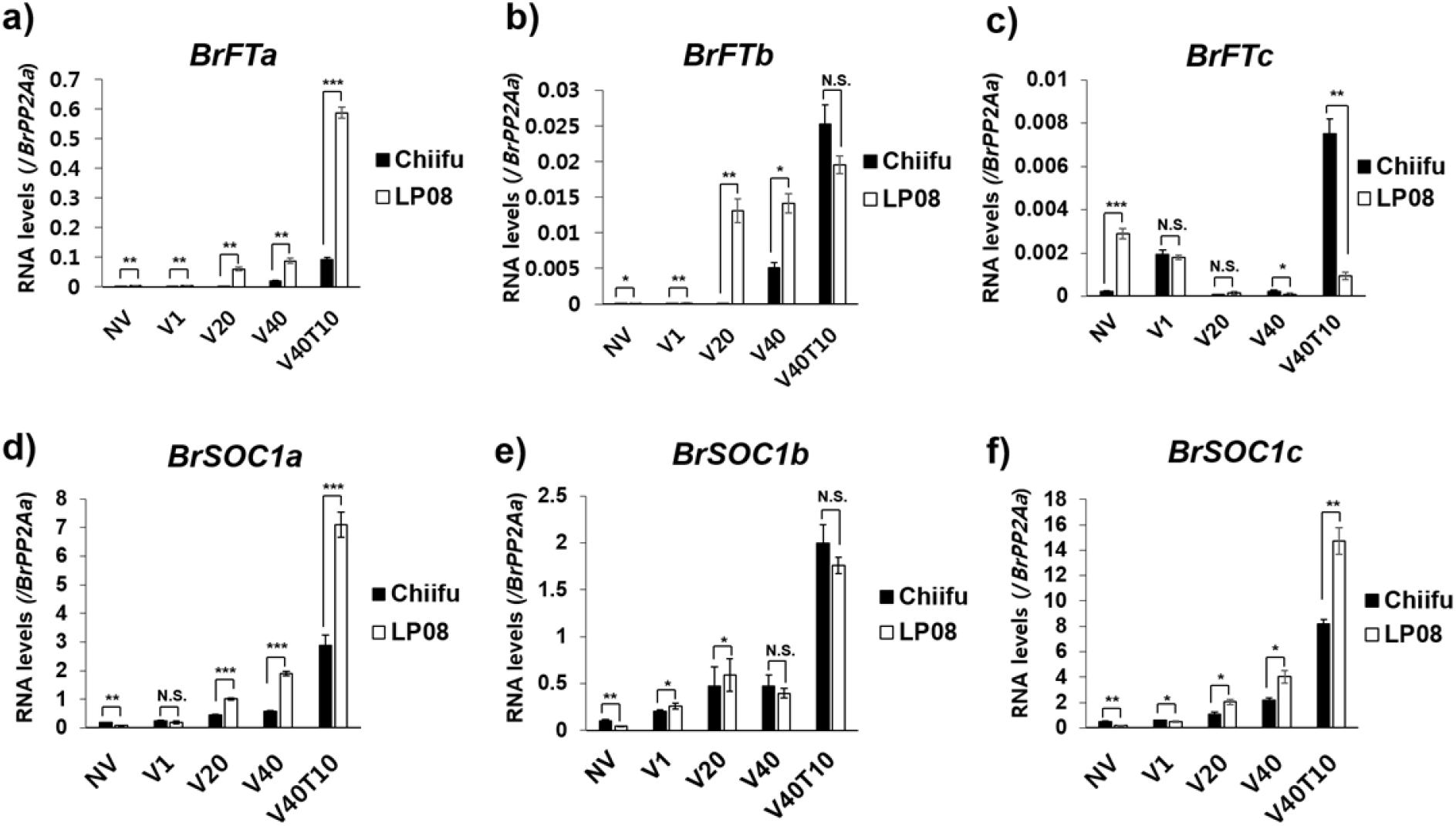
Transcript levels of *BrFT* and *BrSOC1* homologs in ‘Chiifu’ and ‘LP08’ along the vernalization timepoints. Results of qRT-PCR of the three *BrFT* and three *BrSOC1* homologs, (**a**) *BrFTa*, (**b**) *BrFTb*, (**c**) *BrFTc*, (**d**) *BrSOC1a*, (**e**) *BrSOC1b*, and (**f**) *BrSOC1c*, in ‘Chiifu’ and ‘LP08’ along the vernalization time points. *BrPP2Aa* was used as the reference gene. Average and standard errors were calculated using the Ct values of three biological replicates. Significance was statistically determined by t-test (\**p* > 0.05; \*\**p* < 0.01; \*\*\**p* < 0.001).

### Transcription of *FT* and *SOC1* homologs in ‘Chiifu’ and ‘LP08’

We identified the following *FT* and *SOC1* homologs in *B. rapa: BrFTa* (Bra022475), *BrFTb* (Bra004117), *BrFTc* (Bra010052), *BrSOC1a* (Bra000393), *BrSOC1b* (Bra004928), and *BrSOC1c* (Bra039324). The results of qRT-PCR are shown in Figure 5 and Supplementary Figure S4. All homologs were greatly induced upon vernalization, indicating that *BrFT* and *BrSOC1* homologs positively regulate floral transition in Chinese cabbage plants.

The homologs *BrFTa* and, to a lesser extent, *BrFTb* were induced (V40T10), while *BrFTc* was moderately expressed upon vernalization (Fig. 5a–5c and Supplementary Fig. S4a), suggesting that *BrFTa* and *BrFTb* play a role in the induction of floral transition in *B. rapa* plants. In the early flowering line ‘LP08,’ the levels of *BrFTa* and *BrFTb* transcripts were significantly greater than those in ‘Chiifu’ after V20 treatment (Fig. 5a and 5b). Thus, it is likely that early and increased expression of *BrFTa* and *BrFTb* resulted in rapid flowering in ‘LP08’ compared with that in ‘Chiifu.’

Among the three *BrSOC1* homologs, *BrSOC1a* and *BrSOC1c* showed higher expression in ‘LP08’ than in ‘Chiifu,’ whereas, *BrSOC1b* did not exhibit significant difference in transcript levels between the two lines throughout the vernalization treatment (Fig. 5d–5f). This indicated that *BrFLC2* primarily controls the expression of *BrSOC1a* and *BrSOC1c*, and other *BrFLC* homologs (i.e., *BrFLC1* and *BrFLC3)* might play a role in the suppression of *BrSOC1b* in *B. rapa* plants. However, functional redundancy among *BrFLC* homologs in terms of regulation of its downstream targets (*BrFTs* and *BrSOC1s*) in *B. rapa* plants warrants investigation. Moreover, it is noteworthy that all three *BrSOC1* homologs did not exactly match with the predicted gene structure (Supplementary Fig. S4b), wherein the transcript reads corresponding to the first exon of all *BrSOC1* were mapped only to the promoter region, suggesting that the present structures of *BrSOC1* homologs need correction.

### Active degradation of *BrFLC2* in ‘LP08’ by NMD

Integrated QTL mapping and transcriptome analysis of the lines ‘Chiifu’ and ‘LP08’ indicated that *BrFLC2* is the causative gene for rapid flowering in ‘LP08.’ Therefore, to understand the mechanism *BrFLC2* regulation in ‘LP08,’ genomic *BrFLC2* sequences from both ‘Chiifu’ and ‘LP08’ lines were amplified. However, smaller DNA fragments (approximately 6.4 kb) were observed in ‘LP08’ than those in ‘Chiifu’ (approximately 6.9 kb) (Fig. 6a and Supplementary Fig. S5). Additionally, we found that the line ‘LP08’ contained two large deletions (56 bp and 470 bp in sizes), seven single-base substitutions, and two single-base deletions throughout the genomic sequence of *BrFLC2*.

**Figure 6.**
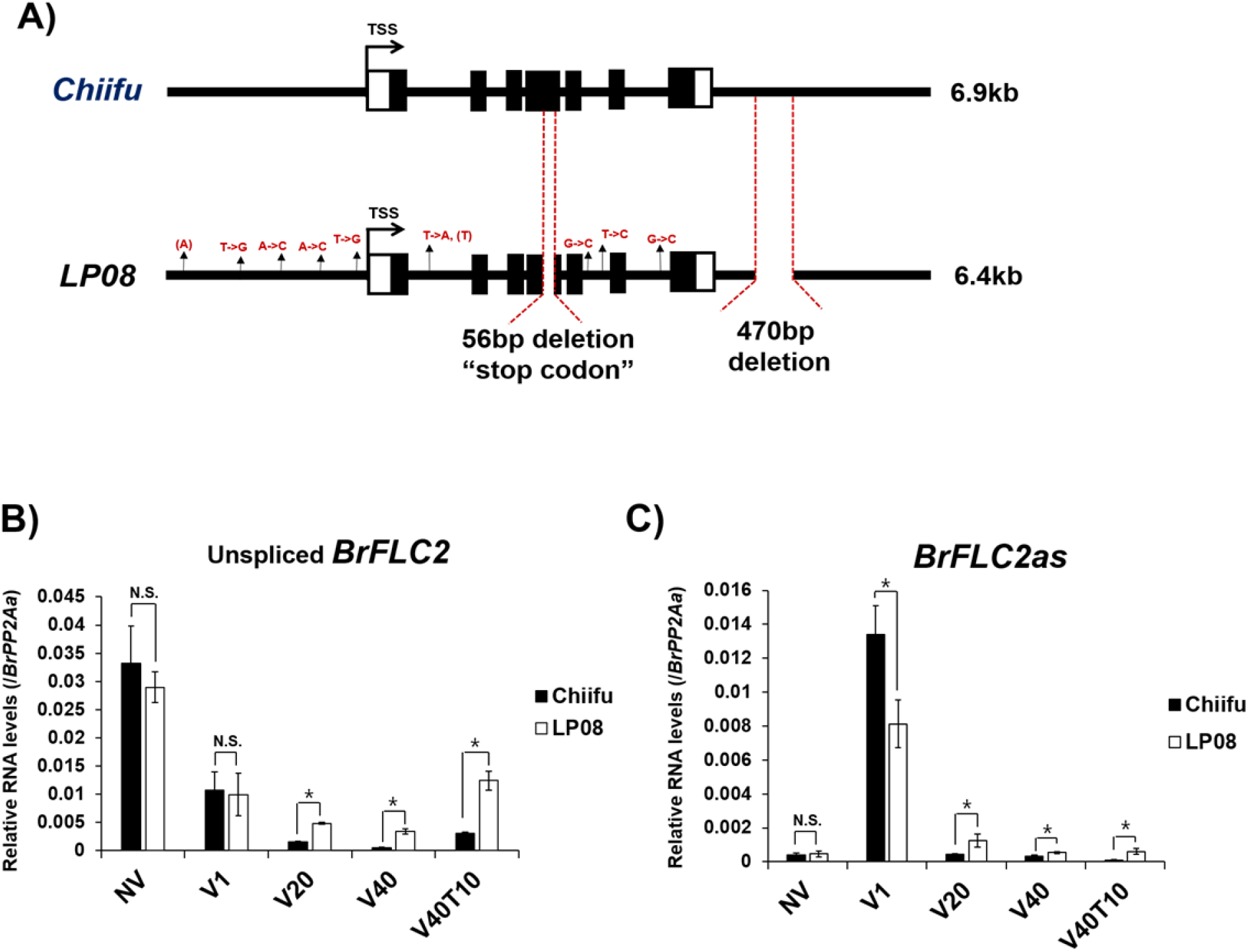
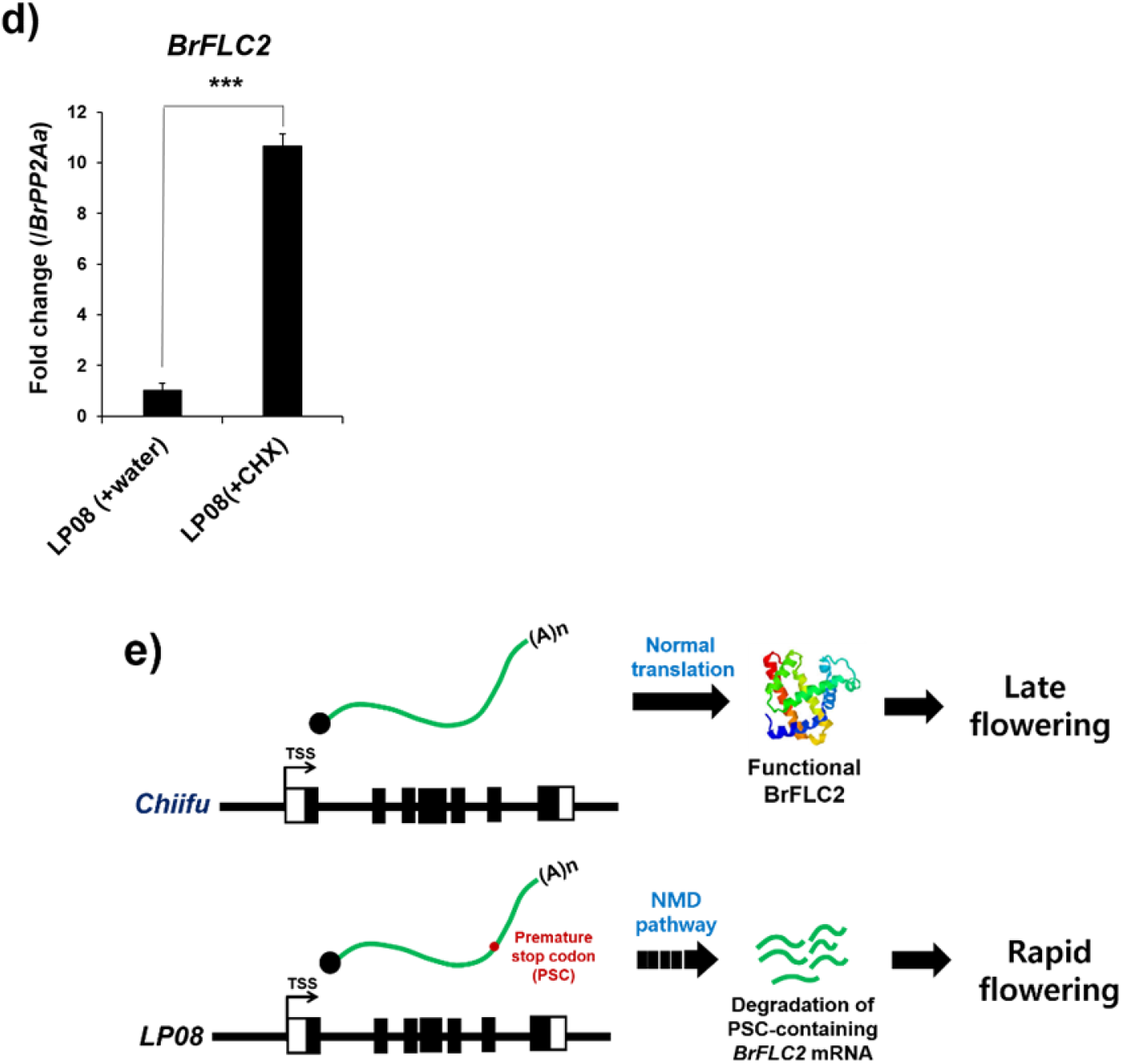
NMD of *BrFLC2* transcripts in ‘LP08.’ (**a**) Comparison of genomic structure of *BrFLC2* between ‘Chiifu’ (upper) and ‘LP08’ (lower). Nine point mutations, two large deletions including 56-bp deletion at the fourth exon-intron junction and 470-bp deletion at the 3’ downstream region were identified in ‘LP08.’ (**b**) Levels of unspliced (nascent) *BrFLC2* transcripts in ‘Chiifu’ and ‘LP08’ along the vernalization time course. (**c**) Levels of antisense ncRNAs, *BrFLC2as*, transcripts in ‘Chiifu’ and ‘LP08’ along the vernalization time course. (**d**) Quantification of *BrFLC2* mRNA levels in control (water-treated) and cycloheximide (CHX)-treated ‘LP08’ plants. Inhibition of NMD by CHX treatment increased the level of BrFLC2 mRNA transcripts. (**e**) Schematic illustration of NMD of *BrFLC2* transcripts. 56-bp deletion on the fourth exon-intron junction produced a premature stop codon (PSC). PSC-containing *BrFLC2* mRNA in ‘LP08’ is downregulated and results in early flowering. Meanwhile, wildtype *BrFLC2* mRNA in ‘Chiifu’ is translated to produce functional BrFLC2, which acts as a floral repressor, conferring late flowering in Chinese cabbage.

We further investigated whether these mutations affected the transcriptional activity of *BrFLC2* in ‘LP08’ using qRT-PCR. First, four single-base substitutions and one single-base deletion were identified in the promoter region of *BrFLC2* in ‘LP08.’ Since some mutations in the promoter region affect transcriptional activity, we measured the levels of nascent RNA transcripts of *BrFLC2* and found that the transcript levels were similar (NV and V1 time points) or higher (V20–V40T10) in ‘LP08’ than in ‘Chiifu’ (Fig. 6b). This suggests that the point mutations in the promoter region of *BrFLC2* in ‘LP08’ did not negatively influence its transcriptional activity. Second, four single-base substitutions and one single-base deletion were identified in the coding region of *BrFLC2* in ‘LP08’ (Fig. 6a),but they were located in the intronic regions destined to be spliced to produce mature mRNA. However, no mutations that could affect splicing were identified at the exon-intron junctions in ‘LP08,’ indicating that the intronic point mutations do not affect the stability of *BrFLC2* mRNA in ‘LP08’.

Next, we investigated the effect of the two large deletions on *BrFLC2* transcription. The 56-bp deletion was found in the fourth exon and the 470-bp deletion was found near the 3’ downstream region (Fig. 6c). Previous reports suggest that *BrFLC2* contains a series of antisense non-coding RNAs (*BrFLC2as*), which are derived from the 3’ downstream region of the sense *BrFLC2* strand (Li et al. 2016), similar to the *COOLAIR* antisense ncRNAs found in *Arabidopsis* (Swiezewski et al. 2009). Moreover, ectopic expression of *BrFLC2as* ncRNA in *B. rapa* plants has been shown to exert a negative effect on *BrFLC2* transcripts. Thus, we examined whether the expression of *BrFLC2as* ncRNA was affected by the 470-bp deletion in ‘LP08.’ Similar to *Arabidopsis COOLAIR* ncRNAs, the transcript levels of *BrFLC2as* were notably increased at V1 and reduced for V20–V40T10 treatments (Fig. 6c). This dynamic pattern was observed in both ‘Chiifu’ and ‘LP08’, indicating that the transcription of *BrFLC2as* ncRNA in ‘LP08’ is not affected by the 470-bp deletion (Fig. 6c).

We then investigated the effect of the 56-bp deletion present in the fourth exon of *BrFLC2* and identified a premature stop codon (PSC), which resulted in a 68-amino acid shorter (at the C-terminus) BrFLC2 (128 amino acids) in ‘LP08’ (Supplementary Figs. S6 and S8). In contrast, full-length BrFLC2 (196 amino acids) in ‘Chiifu’ exhibited high similarities with the FLC homologs from other members of *Brassicaceae* (Supplementary Fig. S8). However, it has been reported that transcripts containing PSCs can be eliminated by a transcript surveillance system referred to as NMD (Brogna and Wen 2009; Schweingruber et al. 2013), which exists in eukaryotes to reduce errors in gene expression.

Since *BrFLC2* transcript in ‘LP08’ contains a PSC, we tested for NMD of *BrFLC2* transcripts in ‘LP08.’ To inhibit NMD, cycloheximide (CHX)-treated seedlings of ‘LP08’ were used. We observed that *BrFLC2* was significantly upregulated in CHX-treated samples compared with that in the control samples (Fig. 6d). Based on these findings, we conclude that low levels of *BrFLC2* in ‘LP08’ can be attributed to NMD, which eliminates abnormal transcripts containing a PSC, resulting in the rapid flowering phenotype in ‘LP08’ (Fig. 6e).

## DISCUSSION

We report that oilseed-type *B. rapa* line ‘LP08’ exhibits earlier flowering than the vegetable-type *B. rapa* ‘Chiifu’ regardless of vernalization treatment. Therefore, to identify the key players responsible for early flowering in ‘LP08,’ we developed a mapping population using the late-flowering line ‘Chiifu’ and the early-flowering line ‘LP08.’ A genetic linkage map generated using 7,833 SNPs was used to pinpoint QTL peaks controlling flowering behavior in the two lines. Since the QTL peak with the maximum LOD score was detected on the upper arm of chromosome A02 (A02:1950553–A02:7744515) in both NV and vernalized conditions, we concluded that this region contained the key regulators contributing to the different flowering behaviors of the two lines.

We also performed comparative transcriptome analysis to identify the transcriptional differences in the candidate floral transition genes determined from QTL mapping. We successfully identified the key player *BrFLC2*, which is responsible for rapid flowering in oilseed-type ‘LP08,’ using integrated QTL mapping and transcriptome analysis. In addition, we found that *BrFLC2* contains a 56-bp deletion in the fourth exon, which generates a PSC, resulting in a truncated BrFLC2. Abnormal *BrFLC2* transcripts in ‘LP08’ are one of the targets of NMD. Previous reports suggest that *BrFLC1* also plays a role in controlling the flowering behavior in *B. rapa* plants, similar to its homolog in Arabidopsis (Wu et al. 2012; Yuan et al. 2009). Thus, we quantified the transcript levels of the four *FLC* homologs in ‘Chiifu’ and ‘LP08’ throughout the vernalization treatment (Fig. 3 and Supplementary Fig. S2). Besides *BrFLC2, BrFLC1* was adequately expressed under NV condition, suggesting that *BrFLC1* also functions as a floral repressor before vernalization. The transcript levels of *BrFLC1* were lower under NV and increased after long-term cold treatments (V20–V40T10) compared with the levels of *BrFLC1* in ‘Chiifu.’ Based on these results, we hypothesized that the lower expression of *BrFLC1* might contribute to rapid flowering of ‘LP08’ under NV condition, but not after vernalization. Thereafter, we detected a significant peak on Chromosome A10 in plants under NV condition (Fig. 1c), where *BrFLC1* (Bra009055) was located, suggesting that *BrFLC1* and *BrFLC2* also affect flowering time in *B. rapa*, particularly under NV condition, which is in agreement with previous reports (Li et al. 2009; Yuan et al. 2009).

Additionally, while flowering time of ‘LP08’ was rapid compared to that of ‘Chiifu,’ the ‘LP08’ line responded to vernalization treatment, i.e., ‘LP08’ flowered earlier after vernalization than under NV condition (Fig. 1a). Although *BrFLC* homologs (*BrFLC1*, *BrFLC3*, and *BrFLC5*), except *BrFLC2*, were sufficiently expressed in ‘LP08’ under NV condition, delayed flowering was observed in ‘LP08’ before vernalization (Fig. 3 and Supplementary Fig. S2), indicating that BrFLC1, BrFLC3, and BrFLC5 play roles as floral repressors in the ‘LP08’ line.

The transcript level of *Arabidopsis FLC* is gradually and stably repressed upon vernalization, wherein VIN3-containing histone modifying complexes (for example, polycomb repressive complex 2) play a pivotal role in repression of *FLC* via histone modifications (Kim and Sung 2013). Recently, it has been reported that a DNA motif, called as COLD MEMORY ELEMENT (CME) located in the first intron of *FLC*, plays an important role in the recruitment of VIN3-containing polycomb complexes in *Arabidopsis* plants (Yuan et al., 2016 Nature Genetics; Questa et al., 2016). Therefore, further investigation is necessary to determine whether the four *BrFLC* homologs *(BrFLC1, BrFLC2, BrFLC3*, and *BrFLC5*) possess CME-like DNA elements in their genomic regions, which might recruit BrVIN3-containing polycomb complexes in *B. rapa* plants.

## MATERIALS AND METHODS

### Plant material and growth conditions

Late-flowering Chinese cabbage (*B. rapa* L. ssp. *pekinensis*) inbred line ‘Chiifu’ and early-flowering, oilseed-type yellow sarson (*B. rapa* L. ssp. *trilocularis)* line ‘LP08’ (S4 generation plant of YS-033, CGN06835) were used in this study. The mapping population comprised 151 F_5_ RILs derived from selfing single F_2_ progeny of the cross between ‘Chiifu’ and ‘LP08’ in the greenhouse of Rural Development Administration of Korea as described previously (Kim et al. 2017). This population was used for QTL mapping of flowering time.

### Construction of genotyping-by-sequencing (GBS) libraries

A total of 153 samples (151 F_5_ plants and 2 parental plants) were subjected to GBS using the Illumina NextSeq500 sequencing workflow (eGnome, South Korea). Genomic DNA was extracted from 0.1g of leaf tissue of each plant using DNeasy Plant Mini Kit (Qiagen, Germany), following the manufacturer’s instructions. The extracted DNA was quantified using Quant-iT PicoGreen dsDNA Assay Kit (Molecular Probes, USA) on a Synergy HTX Multi-Mode Reader (Biotek, USA) and normalized to 20 ng μL^-1^. DNA (200 ng) was digested with 8 U of high-fidelity *Apek*I (New England Biolabs, USA) at 75 °C for 2 h.

DNA libraries for GBS were constructed according to the protocols described previously (Elshire et al. 2011; De Donato et al. 2013), with minor modifications, wherein the restriction digestion of DNA with *Pst* I was followed by adapter ligation. Cohesive-end ligation was performed using a set of 96 different barcode-containing adapters to tag individual samples, a common adapter for all the samples, and 200 U of T4 DNA ligase (New England Biolabs, USA) at 22 °C for 2 h, which was subjected to 65 °C for 20 min for inactivation. Sets of 95 ligations were pooled into one sample and purified using QIAquick PCR Purification Kit (Qiagen, Germany). The pooled ligation mixtures (5 μL) of each sample were amplified using multiplex PCR with AccuPower *Pfu* PCR PreMix (Bioneer, South Korea) and 25 pmol of each primer, making a total volume of 50 μL. PCR cycles consisted of an initial denaturation step at 98 °C for 5 min, followed by 18 cycles of 98 °C for 10 s (denaturation), 65 °C for 5 s (annealing), and 72 °C for 5 s (extension), and a final extension step at 72 °C for 5 min. The PCR products were also purified using QIAquick PCR Purification Kit (Qiagen, Germany) and the fragment sizes were evaluated using 2100 Agilent BioAnalyzer (Agilent Technologies, USA). Thereafter, GBS libraries were sequenced on the Illumina NextSeq500 (Illumina, USA) and single-end reads of length 150 bp were obtained.

### Sequencing and genotyping

Raw data was demultiplexed to sort samples using the GBSX tool (Herten et al. 2015), and chromosome pseudomolecule level genome data (Brassica rapa V2.1 Assembly, BrapaV2.1PacBio.Chr.fa.gz) from *Brassica* database (BRAD) (http://brassicadb.org) was used as the reference. After demultiplexing, single-end reads were mapped to the *B. rapa* reference genome using Bowtie2 (Langmead and Salzberg 2012). For calling variants, we used the software packages Genome Analysis Toolkit (GATK) and Picard tools (McKenna et al. 2010). We also performed local realignment of reads to correct misalignment caused by the presence of indels using GATK ‘RealignerTargetCreator’ and ‘IndelRealigner’ sequence data processing tools. Subsequently, GATK ‘HaplotypeCaller’ and ‘SelectVariants’ tools were used for variant calling.

### Linkage map construction & QTL mapping

Custom code was used to transform variant call format SNP data into input format for R/Qtl package (Broman et al. 2003). Markers with duplicated pattern or distorted segregation ratio, estimated using Chi-square test with *p*-value determined using Bonferroni correction, were filtered. The est.map function of R/Qtl was used to construct the linkage map, along with Kosambi map function to convert genetic distances into recombination fractions, *p*-value threshold of 1e-06, and EM iterations of 1000. Subsequently, the composite interval mapping function of R/Qtl package was used for QTL mapping using Kosambi function. Regions inferred from peak positions with local maximum LOD values, which exceeded the threshold determined using 1000 permutation tests, were selected as significant QTL.

### Measurement of flowering time

Surface-sterilized seeds from parental lines and RILs were grown on half Murashige and Skoog (MS) media at 4 °C for 3 d in dark for stratification. To obtain flowering time data for QTL mapping, 2-week-old seedlings of the 2 parental and 151 F_2_ plants were transplanted into soil pots, transferred to the greenhouse, and then their DTF were counted. Another set of two-week-old seedlings were transferred to soil pots and then moved to the greenhouse for flowering time measurement of the individual lines after 40-day cold treatment (4°C).

### Comparison of genomic sequences of *BrFLC2* of ‘Chiifu’ and ‘LP08’

To compare the genomic sequences of *BrFLC2*, genomic DNA was extracted from ‘Chiifu’ and ‘LP08’ leaves using the TIANamp Genomic DNA kit (Tiangen Biotech, China). Approximately 7 kb-long *BrFLC2* sequence (including the 2 kb promoter, 4 kb coding sequence, and 1 kb 3’ downstream region) from the two lines was PCR-amplified using Solg™ Pfu-X Taq DNA polymerase (SolGent, Republic of Korea). The oligonucleotides were designed based on the sequence information obtained from BRAD (http://brassicadb.org/brad/index.php), and detailed primer information is given in Supplementary Table S4. PCR was performed as follows: the template was initially denatured at 94 °C for 4 min, followed by 38 cycles of amplification (94 °C for 30 s and 68 °C for 8 min), and a final extension at 72 °C for 10 min. The PCR products were purified using Dyne PCR Purification Kit (DyneBio, South Korea) and cloned into pPZP211 vectors using Takara T4 DNA Ligase (Takara Bio, Japan). *Escherichia coli* (DH5α) cells were transformed with the plasmid DNA carrying the *BrFLC2* PCR product.

### RNA extraction and qRT-PCR analysis

Total RNA was extracted from whole seedling samples using RNeasy Mini Kit (Qiagen, Germany) and treated with DNase I (New England Biolabs, USA) to remove genomic DNA contamination. qRT-PCR was performed using Solg™ 2X Real-time PCR Smart Mix (SolGent, South Korea) under the following cycling conditions: initial denaturation at 95 °C for 12 min, followed by 50 cycles at 95 °C for 20 s, 60 °C for 25 s, and 72 °C 35 s. The primer sequences were designed based on sequence information from BRAD (Supplementary Table S3). *BrPP2Aa* (Bra012474), which exhibited a consistent expression at each vernalization time point in our RNA-seq data, was used as the reference gene. Three biological replicates were analyzed for each qRT-PCR assay, and Student’s *t*-test was used for statistical analyses.

### RNA-seq library construction and sequencing

Three biological replicates for each time point were harvested and frozen in liquid nitrogen. Total RNA was extracted, and RNA-seq libraries were constructed using TruSeq Stranded mRNA LT Sample Prep Kit (Illumina Inc., USA) according to the manufacturer’s instructions. Libraries were then sequenced on NovaSeq 6000 system (Macrogen Inc., South Korea) using the paired-end sequencing protocol.

### RNA-seq alignment and analysis

Quality of the RNA-seq reads were evaluated using the FastQC software (http://www.bioinformatics.babraham.ac.uk/projects/fastqc). The raw reads were trimmed and quality-filtered before alignment, and those with more than 90 % threshold (Q>30) were used for mapping. *B. rapa* reference genome was obtained from Ensembl (https://plants.ensembl.org/info/website/ftp/index.html), and mapping was performed using the TopHat2 software with default parameters (Kim et al. 2013). Aligned reads were converted to digital counts using HTseq-count and were analyzed using edgeR. Differentially expressed genes (DEGs) were identified based on a 0.05 *p*-value and a cutoff of two-fold difference in expression. Multi-dimensional scaling plot and correlation heap map were generated using R packages (ver. 3.6.0). Venn diagram and GO analyses were performed using Venny webtool (ver. 2.1) (https://bioinfogp.cnb.csic.es/tools/venny/) and ShinyGO (ver 0.61) program (Ge et al. 2020), respectively. Aligned reads were then converted to bigwig files for visualization using Integrative Genomics Viewer program of the Broad Institute (Thorvaldsdottir et al. 2013).

### NMD assay

Seeds of ‘LP08’ were sterilized and plated on half MS medium containing 2.5 % sucrose and 0.6 % plant agar (Duchefa Biochemia, Netherland). After three d of stratification, seedlings were grown in a growth room at 22 °C and 16-h light/8-h dark photoperiod for 7 d. For CHX (Sigma-Aldrich, USA) treatment, seedlings were dipped in half MS liquid medium containing 2 % sucrose and 10 μg mL^-1^ CHX or water (control) (Rayson et al. 2012), incubated for 4 h at RT with slight shaking, and then frozen for RNA extraction. Total RNA was extracted and treated with DNaseI. First-strand cDNA was synthesized using 5 μg of total RNA and EasyScript Reverse Transcriptase (TransGen, China). The cDNA was diluted using distilled water and used for qRT-PCR reaction.

## ACKNOWLEDGEMENTS

This work was supported by grants from the BioGreen 21 Agri-Tech Innovation Program of the Rural Development Administration, Republic of Korea (project No. PJ01566203) to D. H. K.

## AUTHORS CONTRIBUTIONS

S.K., J.A.K., and H.K. derived the RILs and performed the molecular experiments; S.K., J.A.K., and D.H.K. planned the experiments and analyzed the data; D.H.K. wrote the manuscript.

## Parsed Citations

Amasino R (2010) Seasonal and developmental timing of flowering. Plant J 61 (6):1001–1013. doi:10.1111/j.1365-313X.2010.04148.x

Bastow R, Mylne JS, Lister C, Lippman Z, Martienssen RA, Dean C (2004) Vernalization requires epigenetic silencing of FLC by histone methylation. Nature 427 (6970):164–167. doi:10.1038/nature02269

Brogna S, Wen JK (2009) Nonsense-mediated mRNA decay (NMD) mechanisms. Nat Struct Mol Biol 16 (2):107–113. doi:10.1038/nsmb.1550

Broman KW, Wu H, Sen S, Churchill GA (2003) R/qtl: QTL mapping in experimental crosses. Bioinformatics 19 (7):889–890. doi:10.1093/bioinformatics/btg112

De Donato M, Peters SO, Mitchell SE, Hussain T, Imumorin IG (2013) Genotyping-by-Sequencing (GBS): A Novel, Efficient and Cost-Effective Genotyping Method for Cattle Using Next-Generation Sequencing. Plos One 8 (5). doi: ARTN e62137

Dennis ES, Helliwell CA, Peacock WJ (2006) Vernalization: Spring into flowering. Dev Cell 11 (1):1–2. doi: 10.1016/j.devcel.2006.06.007 10.1371/journal.pone.0062137

Elshire RJ, Glaubitz JC, Sun Q, Poland JA, Kawamoto K, Buckler ES, Mitchell SE (2011) A Robust, Simple Genotyping-by-Sequencing (GBS) Approach for High Diversity Species. Plos One 6 (5). doi: ARTN e19379

Ge SX, Jung DM, Yao RA (2020) ShinyGO: a graphical gene-set enrichment tool for animals and plants. Bioinformatics 36 (8):2628–2629. doi: 10.1093/bioinformatics/btz931 10.1371/journal.pone.0019379

Geraldo N, Baurle I, Kidou S, Hu XY, Dean C (2009) FRIGIDA Delays Flowering in Arabidopsis via a Cotranscriptional Mechanism Involving Direct Interaction with the Nuclear Cap-Binding Complex. Plant Physiology 150 (3):1611–1618. doi: 10.1104/pp.109.137448

Herten K, Hestand MS, Vermeesch JR, Van Houdt JK (2015) GBSX: a toolkit for experimental design and demultiplexing genotyping by sequencing experiments. BMC bioinformatics 16 (1):1

Kawanabe T, Osabe K, Itabashi E, Okazaki K, Dennis ES, Fujimoto R (2016) Development of primer sets that can verify the enrichment of histone modifications, and their application to examining vernalization-mediated chromatin changes in Brassica rapa L. Genes Genet Syst 91 (1):1–10. doi: 10.1266/ggs.15-00058

Kim D, Pertea G, Trapnell C, Pimentel H, Kelley R, Salzberg SL (2013) TopHat2: accurate alignment of transcriptomes in the presence of insertions, deletions and gene fusions. Genome Biol 14 (4). doi: ARTN R36 10.1186/gb-2013-14-4-r36

Kim DH (2020) Current understanding of flowering pathways in plants: focusing on the vernalization pathway in Arabidopsis and several vegetable crop plants. Hortic Environ Biote 61 (2):209–227. doi: 10.1007/s13580-019-00218-5

Kim DH, Doyle MR, Sung S, Amasino RM (2009) Vernalization: Winter and the Timing of Flowering in Plants. Annu Rev Cell Dev Bi 25:277–299. doi: 10.1146/annurev.cellbio.042308.113411

Kim DH, Sung S (2013) Coordination of the vernalization response through a VIN3 and FLC gene family regulatory network in Arabidopsis. Plant Cell 25 (2):454–469. doi: 10.1105/tpc.112.104760

Kitamoto N, Yui S, Nishikawa K, Takahata Y, Yokoi S (2014) A naturally occurring long insertion in the first intron in the Brassica rapa FLC2 gene causes delayed bolting. Euphytica 196 (2):213–223. doi: 10.1007/s10681-013-1025-9

Koornneef M, Blankestijndevries H, Hanhart C, Soppe W, Peeters T (1994) The Phenotype of Some Late-Flowering Mutants Is Enhanced by a Locus on Chromosome-5 That Is Not Effective in the Landsberg Erecta Wild-Type. Plant Journal 6 (6):911–919. doi: DOI 10.1046/j.1365-313X.1994.6060911.x

Langmead B, Salzberg SL (2012) Fast gapped-read alignment with Bowtie 2. Nat Methods 9 (4):357–U354. doi: 10.1038/Nmeth.1923

Lee I, Michaels SD, Masshardt AS, Amasino RM (1994) The Late-Flowering Phenotype of Frigida and Mutations in Luminidependens Is Suppressed in the Landsberg Erecta Strain of Arabidopsis. Plant Journal 6 (6):903–909. doi: DOI 10.1046/j.1365-313X.1994.6060903.x

Leijten W, Koes R, Roobeek I, Frugis G (2018) Translating Flowering Time from Arabidopsis thaliana to Brassicaceae and Asteraceae Crop Species. Plants-Basel 7 (4). doi: ARTN 111 10.3390/plants7040111

Li F, Kitashiba H, Inaba K, Nishio T (2009) A Brassica rapa Linkage Map of EST-based SNP Markers for Identification of Candidate Genes Controlling Flowering Time and Leaf Morphological Traits. DNA Res 16 (6):311–323. doi: 10.1093/dnares/dsp020

Li XR, Zhang SF, Bai JJ, He YK (2016) Tuning growth cycles of Brassica crops via natural antisense transcripts of BrFLC. Plant Biotechnol J 14 (3):905–914. doi: 10.1111/pbi.12443

McKenna A, Hanna M, Banks E, Sivachenko A, Cibulskis K, Kernytsky A, Garimella K, Altshuler D, Gabriel S, Daly M (2010) The Genome Analysis Toolkit: a MapReduce framework for analyzing next-generation DNA sequencing data. Genome research 20 (9):1297–1303

Osborn TC, Kole C, Parkin IAP, Sharpe AG, Kuiper M, Lydiate DJ, Trick M (1997) Comparison of flowering time genes in Brassica rapa, B-napus and Arabidopsis thaliana. Genetics 146 (3):1123–1129

Rayson S, Arciga-Reyes L, Wootton L, Zabala MD, Truman W, Graham N, Grant M, Davies B (2012) A Role for Nonsense-Mediated mRNA Decay in Plants: Pathogen Responses Are Induced in Arabidopsis thaliana NMD Mutants. Plos One 7 (2). doi: ARTN e31917

Schweingruber C, Rufener SC, Zund D, Yamashita A, Muhlemann O (2013) Nonsense-mediated mRNA decay - Mechanisms of substrate mRNA recognition and degradation in mammalian cells. Bba-Gene Regul Mech 1829 (6-7):612–623. doi: 10.1016/j.bbagrm2013.02.005 10.1371/journal.pone.0031917

Searle I, He YH, Turck F, Vincent C, Fornara F, Krober S, Amasino RA, Coupland G (2006) The transcription factor FLC confers a flowering response to vernalization by repressing meristem competence and systemic signaling in Arabidopsis. Gene Dev 20 (7):898–912. doi: 10.1101/gad.373506

Sung S, Amasino RM (2005) Remembering winter: Toward a molecular understanding of vernalization. Annu Rev Plant Biol 56:491–508. doi: 10.1146/annurev.arplant.56.032604.144307

Sung SB, Amasino RM (2004) Vernalization in Arabidopsis thaliana is mediated by the PHD finger protein V1N3. Nature 427 (6970):159–164. doi: 10.1038/nature02195

Swiezewski S, Liu FQ, Magusin A, Dean C (2009) Cold-induced silencing by long antisense transcripts of an Arabidopsis Polycomb target. Nature 462 (7274):799–U122. doi: 10.1038/nature08618

Teutonico RA, Osborn TC (1995) Mapping loci controlling vernalization requirement in Brassica rapa. Theor Appl Genet 91 (8):1279–1283. doi: Doi 10.1007/Bf00220941

Thorvaldsdottir H, Robinson JT, Mesirov JP (2013) Integrative Genomics Viewer (IGV): high-performance genomics data visualization and exploration. Brief Bioinform 14 (2):178–192. doi: 10.1093/bib/bbs017

Wu, J, Wei KY, Cheng F, Li SK, Wang Q, Zhao JJ, Bonnema G, Wang XW (2012) A naturally occurring InDel variation in BraAFLC.b(BrFLC2) associated with flowering time variation in Brassica rapa. Bmc Plant Biol 12. doi: Artn 151

Xiao D, Zhao JJ, Hou XL, Basnet RK, Carpio DPD, Zhang NW, Bucher J, Lin K, Cheng F, Wang XW, Bonnema G (2013) The Brassica rapa FLC homologue FLC2 is a key regulator of flowering time, identified through transcriptional co-expression networks. J Exp Bot 64 (14):4503–4516. doi: 10.1093/jxb/ert264 10.1186/1471-2229-12-151

Yuan YX, Wu J, Sun RF, Zhang XW, Xu DH, Bonnema G, Wang XW (2009) A naturally occurring splicing site mutation in the Brassica rapa FLC1 gene is associated with variation in flowering time. J Exp Bot 60 (4):1299–1308. doi: 10.1093/jxb/erp010

Zhao JJ, Kulkarni V, Liu NN, Del Carpio DP, Bucher J, Bonnema G (2010) BrFLC2 (FLOWERING LOCUS C) as a candidate gene for a vernalization response QTL in Brassica rapa. J Exp Bot 61 (6):1817–1825. doi: 10.1093/jxb/erq048

